# SmcHD1 underlies the formation of H3K9me3 blocks on the inactive X chromosome in mice

**DOI:** 10.1101/2021.08.23.457321

**Authors:** Saya Ichihara, Koji Nagao, Takehisa Sakaguchi, Chikashi Obuse, Takashi Sado

## Abstract

Stable silencing of the inactive X chromosome (Xi) in female mammals is critical for the development of embryos and their postnatal health. SmcHD1 is essential for stable silencing of the Xi, and its functional deficiency results in derepression of many X-inactivated genes. Although SmcHD1 has been suggested to play an important role in the formation of higher order chromatin structure of the Xi, the underlying mechanism is largely obscure. Here we explore the epigenetic state of the Xi in SmcHD1-deficient epiblast stem cells (EpiSCs) and mouse embryonic fibroblasts (MEFs) in comparison with their wild-type counterparts. The results suggest that SmcHD1 underlies the formation of H3K9me3-enriched blocks on the Xi, which, although the importance of H3K9me3 has been largely overlooked in mice, play a critical role in the establishment of the stably silenced state. We propose that the H3K9me3 blocks formed on the Xi facilitate robust heterochromatin formation in combination with H3K27me3, and the substantial loss of H3K9me3 caused by SmcHD1 deficiency leads to aberrant distribution of H3K27me3 on the Xi and derepression of X-inactivated genes.

## Introduction

Normal development of organisms requires the proper construction and maintenance of transcriptionally active euchromatin and transcriptionally inactive heterochromatin at different sites in the genome, depending on the developmental stage and cell type. Deposition and removal of epigenetic modifications are thought to be important for these processes. X chromosome inactivation (XCI), a process by which dosage imbalance of X-linked genes between the sexes in mammals is compensated (Lyon, 1961), is a paradigm for understanding how chromatin organization is regulated. In female mice, the paternally derived X chromosome (Xp) becomes transcriptionally silenced at around the 4- to 8-cell stage (Mak *et al*, 2004; Okamoto *et al*, 2004). At the blastocyst stage, however, the Xp that was inactivated early on recovers its transcriptional activity in those cells that have contributed to the inner cell mass (ICM) (Mak *et al*, 2004; Okamoto *et al*, 2004). These ICM cells form a still undifferentiated cell population called the epiblast, giving rise to all the tissues of the fetus after implantation, in which either X chromosome has undergone inactivation regardless of its parental origin.

The noncoding *Xist* gene (Brockdorff *et al*, 1991; Borsani *et al*, 1991) on the X chromosome plays an indispensable role in the initiation of XCI (Penny *et al*, 1996; Marahrens *et al*, 1997). Transcriptional upregulation of *Xist* and the subsequent accumulation of the *Xist* transcript on the X chromosome it originates from are thought to trigger stepwise processes of XCI involving epigenetic modifications such as histone modifications and DNA methylation to form robust heterochromatin (Csankovszki *et al*, 2001; Kohlmaier *et al*, 2004; Wutz & Jaenisch, 2000). Immunofluorescence analyses have demonstrated that the inactive X chromosome (Xi) is enriched in histone H2A monoubiquitinated at lysine 119 (H2AK119ub) and histone H3 trimethylated at lysine 27 (H3K27me3), whose respective post-translational modifications are catalyzed by Polycomb Repressive Complexes PRC1 and PRC2, respectively (Fang *et al*, 2004; Napoles *et al*, 2004; Plath *et al*, 2003; Silva *et al*, 2003; Erhardt *et al*, 2003). Species-specific differences in histone modifications on the Xi, however, have also been pointed out. Antibodies against histone H3 trimethylated at lysine 9 (H3K9me3) and H4 trimethylated at lysine 20 (H4K20me3) highlight the Xi in immunofluorescence assays in marsupials and some eutherians, including humans (Chaumeil *et al*, 2011; Chadwick & Willard, 2004; Rens *et al*, 2010; Zakharova *et al*, 2011), but their enrichments on the Xi are not evident in mice. This would partly explain why most studies of histone modifications in the mechanism of XCI have been focused on those mediated by PRC1 and PRC2 in mice. H3K9me3 and H4K20me3 are known to be enriched at DAPI-dense constitutive heterochromatin such as pericentromeric heterochromatin and telomeres in mammalian somatic cells (Peters *et al*, 2001; Schotta *et al*, 2004). A prevailing view is, therefore, that the mouse Xi is exceptional in that it may not share the mechanism used by the above-noted marsupials and eutherians to establish and maintain the repressed chromatin state with constitutive heterochromatin. A more recent study employing ChIP-seq, however, demonstrated that H3K9me3 was enriched on substantial regions of the Xi compared to the active X chromosome (Xa) in the mouse (Keniry *et al*, 2016).

Mutations of EED, one of PRC2’s core components essential for PRC2-mediated production of H3K27me3, cause derepression of genes on the Xi in the trophoblast of the postimplantation mouse embryo, suggesting that H3K27me3 is not essential for the initiation of XCI, but is important for its maintenance (Kalantry & Magnuson, 2006; Kalantry *et al*, 2006; Montgomery *et al*, 2005; Wang *et al*, 2001). DNA methylation is also known to contribute to sustaining the inactivated state of the Xi (Mohandas *et al*, 1981). CpG islands, which are often found in regulatory regions such as promoters and enhancers, are highly methylated on the Xi as compared to those on the Xa. When cells are treated with DNA demethylating agents such as 5-aza-cytidine, sporadic reactivation of the previously silenced genes on the Xi often takes place in culture. Initiation of random XCI, on the other hand, is not affected in the epiblast lineage of mouse embryos deficient for de novo DNA methyltransferases (Sado *et al*, 2004). These observations suggest that DNA methylation is dispensable for the initiation of XCI but plays a role in the maintenance of XCI. Another factor known to be involved in the maintenance of XCI is SmcHD1 (Blewitt *et al*, 2008). SmcHD1 is a noncanonical member of the SMC family of proteins and accumulates on the Xi. In female mouse embryos homozygous for a null mutation of *Smchd1*, *Smchd1^MommeD1^* (referred to hereafter as *Smchd1^MD1^*), although one of the two X chromosomes undergoes random XCI during early postimplantation development, many genes on the X chromosome thus inactivated become reactivated later on as development progresses, resulting in embryonic lethality at mid-gestation stages (Blewitt *et al*, 2008) Derepression of X-inactivated genes in the homozygous mutant female embryos is accompanied by a reduction in CpG methylation and H3K27me3, suggesting that SmcHD1 is involved in the establishment of the epigenetic state required for the stable maintenance of XCI (Sakakibara *et al*, 2018). More recent studies have also suggested that SmcHD1 is involved in the organization of the higher-order chromatin structure of the Xi (Wang *et al*, 2018; Jansz *et al*, 2018a; Gdula *et al*, 2019), and this is consistent with our previous finding that depletion of SMCHD1diminishes condensed Barr body conformation in human somatic cells (Nozawa *et al*, 2013).

Although our previous study demonstrated the role of SmcHD1 in the establishment of the epigenetic state, it did not address how the aberrant epigenetic state of the Xi resulting in the derepression of X-inactivated genes was created in the absence of SmcHD1, since the findings were essentially based on observations in *Smchd1^MD1/MD1^* mouse embryonic fibroblasts (MEFs) (Sakakibara *et al*, 2018). In this study, we further explored the chromatin state of the Xi and its silencing status using epiblast stem cells (EpiSCs) established from *Smchd1^MD1/MD1^* embryos as a model of the postimplantation epiblast that has just undergone XCI. Many of the derepressed genes on the Xi in SmcHD1-deficient MEFs were repressed with enrichment of H3K27me3 in SmcHD1-deficient EpiSCs, suggesting that genes derepressed on the Xi at later developmental stages acquired H3K27me3 at an early phase of XCI but lost it later on in SmcHD1 mutant embryos. Furthermore, the most prominent feature caused by the lack of SmcHD1 in EpiSCs and MEFs was an extensive loss of H3K9me3 enrichment on the Xi, which apparently disturbed the distribution of H3K27me3. Our findings shed light on the hitherto unexpected role of H3K9me3 in the maintenance of XCI, which has not been extensively studied, especially in the mouse. We discuss the role of SmcHD1 in forming H3K27me3 and H3K9me3 blocks on the Xi and their effects on generating robust heterochromatin.

## Results

### Derivation of EpiSCs from *Smchd1^MD1/MD1^* embryos

It is known that SmcHD1 is essential for stable maintenance of XCI in both the extraembryonic tissues that give rise to the placenta and some extraembryonic membranes, and the embryonic tissue forming the embryo proper. Our previous study suggested that the X-inactivated genes gradually became derepressed during postimplantation development in the absence of SmcHD1 (Sakakibara *et al*, 2018). Allele-resolved RNA-seq further revealed that half of the informative X-linked genes on the Xi were expressed in *Smchd1^MD1/MD1^* MEFs (Sakakibara *et al*, 2018). Although these results suggest that a relatively large fraction of the X-linked genes inactivated early on become derepressed by the mid-gestation stage in the SmcHD1 mutant, the chromosome-wide kinetics of derepression are not clear. To study the XCI state at the early stages of postimplantation development, we took advantage of EpiSCs (Brons *et al*, 2007; Tesar *et al*, 2007) as a model for those cells that have just initiated XCI. EpiSCs have been suggested to resemble postimplantation epiblasts, especially the anterior primitive streak of the epiblast at the late gastrulation-stage, in their cellular and molecular phenotypes (Chen *et al*, 2016; Kojima *et al*, 2014).

We established EpiSCs from the epiblasts of E6.5 embryos recovered from females doubly heterozygous for a dysfunctional *Xist* allele, either *Xist^ΔA^* (Hoki *et al*, 2009) or *Xist^1lox^* (Sado *et al*, 2005), and *Smchd1^MD1^* crossed with males heterozygous for *Smchd1^MD1^* and carrying an X chromosome derived from JF1 (*Mus musculus molosinus*) (X^JF1^) (Figure 1A) (see Materials and methods). Since the dysfunctional *Xist* allele resided on an X chromosome derived from C57Bl/6J (*Mus musculus domesticus*) (X^B6-ΔA^ or X^B6-1lox^), XCI in these EpiSCs had been exclusively skewed to X^JF1^. In addition, the presence of many single nucleotide polymorphisms (SNPs) and insertions/deletions (INDELs) between these subspecies enabled us to carry out allele-resolved analysis of X-linked genes. Each of the established EpiSC lines was genotyped by PCR, and a combination of one wild-type and one homozygous mutant line with respect to the *Smchd1* allele, either [X^B6-ΔA^X^JF1^; *Smchd1^+/+^*] and [X^B6-ΔA^X^JF1^; *Smchd1^MD1/MD1^*] or [X^B6-1lox^X^JF1^; *Smchd1^+/+^*] and [X^B6-1lox^X^JF1^; *Smchd1^MD1/MD1^*], was used for further analyses. RNA-seq described below confirmed that these EpiSCs expressed typical marker genes for EpiSCs (Supplementary Figure S1A).

**Figure 1.**
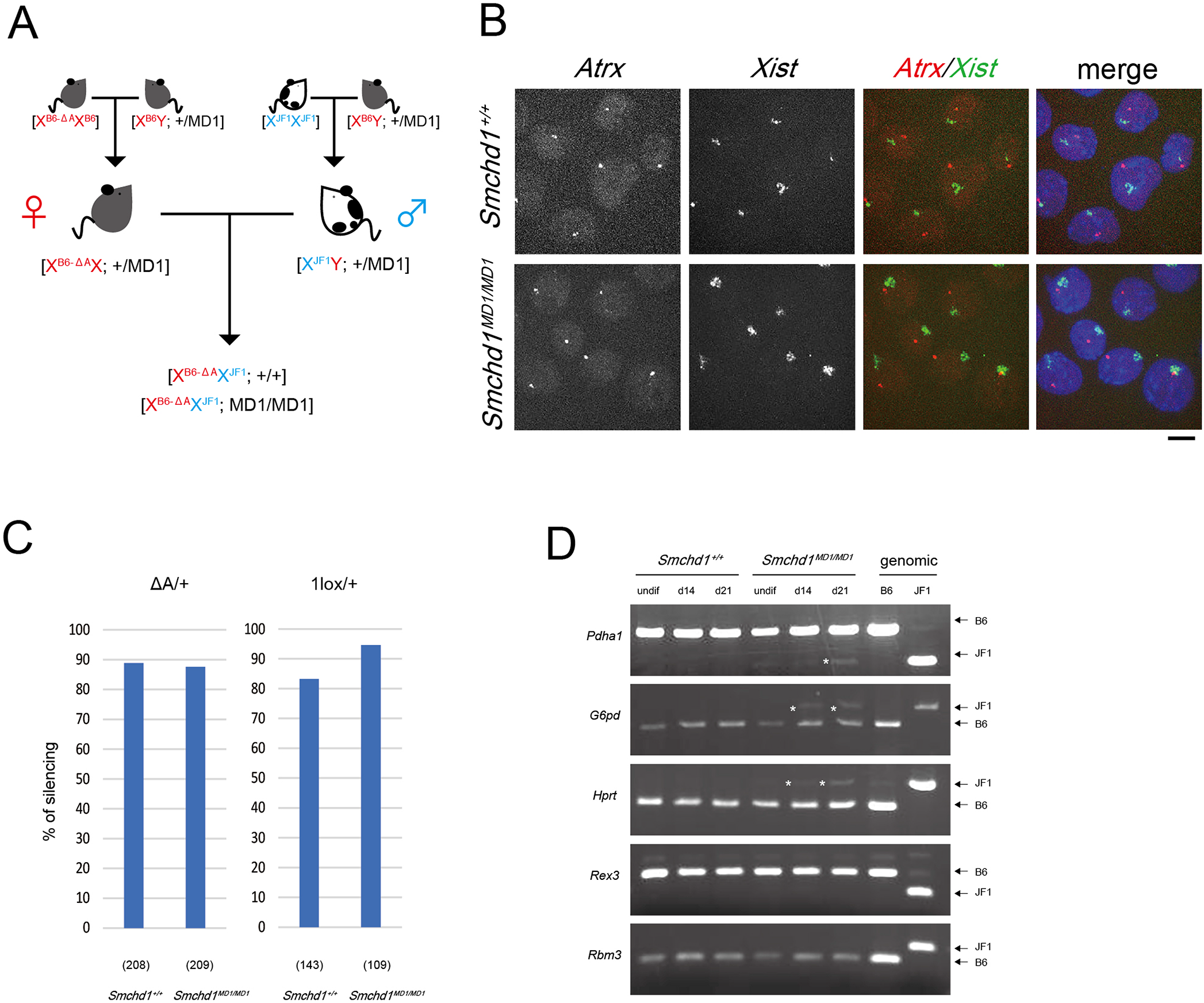
Expression of the X-linked *Atrx* gene in *Smchd1^MD1/MD1^* EpiSCs. (A) Scheme of mouse intercrosses to obtain postimplantation epiblasts carrying respective genotypes, from which EpiSCs were derived. (B) Representative images of RNA-FISH detecting *Xist* (green) and *Atrx* (red) expression in the nuclei of wild-type and *Smchd1^MD1/MD1^* EpiSCs. Scale bar: 10 μm. (C) Proportions of the nuclei with a single pinpoint for *Atrx*, which did not overlap with the *Xist* cloud, in wild-type and *Smchd1^MD1/MD1^* EpiSCs on either *Xist^ΔA^*/+ or *Xist^1lox^*/+ background. The numbers of nuclei examined are shown in parentheses. (D) Allelic expression of X-linked genes in EpiSCs before and after induction of differentiation. A fragment containing a restriction site polymorphism amplified by RT-PCR was digested with an appropriate restriction enzyme and electrophoresed. Digestion of a PCR fragment amplified on genomic DNA of B6 and JF1 is also shown in parallel to indicate allelic difference of the restriction fragment, Expression of *Pdha1*, *G6pd*, and *Hprt* from the JF1 allele was detected only in differentiating (d14 and d21) *Smchd1^MD1/MD1^* EpiSCs. An asterisk highlights a fragment indicative of derepression from the inactive X^JF1^ on the left.

We first examined XCI status in EpiSCs deficient for SmcHD1 by two-color RNA-FISH for *Xist* and X-linked *Atrx* expression. *Atrx* has been shown to be gradually derepressed during postimplantation development in *Smchd1^MD1/MD1^* embryos (Sakakibara *et al*, 2018). In wild-type EpiSCs, expression of *Atrx* was detected as a pinpoint signal, which did not overlap with the *Xist* cloud, in the great majority of cells positive for *Xist* (Figure 1B and 1C), indicating that *Xist* RNA accumulated on the Xi repressed *Atrx*. The situation was essentially the same in *Smchd1^MD1/MD1^* EpiSCs, suggesting that *Atrx* was repressed on the Xi even in the absence of SmcHD1. This contrasted with our previous finding in MEFs prepared from *Smchd1^MD1/MD1^* fetuses at E13.5, in which *Atrx* was derepressed on the Xi coated with *Xist* RNA in 59% of cells (Sakakibara *et al*, 2018).

To study allelic expression of some additional X-linked genes in EpiSCs before and after induction of differentiation, we carried out RT-PCR by amplifying a fragment containing a restriction site polymorphism between JF1 and B6 and subsequent restriction digestion of the fragment. As mentioned earlier, X^JF1^ was exclusively inactivated in the EpiSCs, and therefore, appearance of the JF1-type restriction fragment indicated failure in silencing of the Xi. As shown in Figure 1D, all 5 genes examined were exclusively expressed from the B6 alleles in wild-type EpiSCs not only before but also after differentiation, suggesting stable silencing of genes on the Xi. On the other hand, while these genes were repressed in *Smchd1^MD1/MD1^* EpiSCs before differentiation, *Pdha1*, *G6pd*, and *Hprt* were found to be expressed from the JF alleles in addition to being expressed from the B6 alleles in EpiSCs after induction of differentiation (Figure 1D). The derepression of these genes in differentiating mutant EpiSCs was consistent with the phenotype of *Smchd1*-deficient embryos, in which X-linked genes inactivated early on by the accumulation of *Xist* RNA gradually became derepressed at later stages of development. These results supported the idea that EpiSCs could be used as a model for the early postimplantation-stage epiblast to study the effect of SmcHD1-deficiency at a relatively early phase of XCI.

### The majority of genes on the Xi were repressed in S*mchd1^MD1/MD1^* EpiSCs

To further characterize the X-inactivated state in SmcHD1-deficient EpiSCs in more detail, we performed allele-resolved RNA-seq. Since X^JF1^ was exclusively inactivated in EpiSCs used in this study due to the deficiency of *Xist* on the other B6-derived X, detection of sequence reads carrying JF1-type SNPs or INDELs should be indicative of a failure of repression on the otherwise inactive X^JF1^. We compared Xi-probability of respective X-linked genes (% Xi) [paternal reads/ (maternal reads + paternal reads)] between wild-type and *Smchd1^MD1/MD1^* EpiSCs. Genes with an Xi-probability of 10% or higher were referred to as those expressed from the Xi. The majority of genes in wild-type EpiSCs were scored lower than 10%, indicating that they were essentially inactivated as expected (Figure 2A). Of 387 informative X-linked genes, 18 (4.7%) deviated from this category in wild-type EpiSCs, which was comparable to the number in wild-type MEFs (11/449, 2.4%). They might have not yet been inactivated or might have escaped inactivation in EpiSCs. This was also the case in another line of EpiSCs (Supplementary Figure S1B and S1C). In contrast to wild-type EpiSCs, 21% (77/369) of the X-linked genes in *Smchd1^MD1/MD1^* EpiSCs showed Xi-probability higher than 10%, and therefore were considered to be misexpressed from the inactive X^JF1^ (Figure 2A). Although it was reasonable to assume that such misexpression was caused by the lack of SmcHD1, it was not clear if these genes were derepressed after inactivation at the earlier stages, or if instead they failed to be inactivated from the beginning. Nonetheless, they are referred to as derepressed genes hereafter. Although a fraction of genes on the Xi were derepressed in *Smchd1^MD1/MD1^* EpiSCs, cumulative distribution plots showed that changes in X-linked gene expression between wild-type and SmcHD1-deficient EpiSCs were comparable to the autosomal profile, in contrast to the changes in MEFs, indicating that the Xi genes were not substantially upregulated in *Smchd1^MD1/MD1^* EpiSCs (Supplementary Figure S1D). Figure 2B shows the overlap of the derepressed genes in *Smchd1^MD1/MD1^* MEFs and EpiSCs among the 354 informative genes commonly silenced on the Xi in both wild-type EpiSC lines. Of 72 derepressed genes in *Smchd1^MD1/MD1^* EpiSCs, 62 were included in those derepressed in *Smchd1^MD1/MD1^* MEFs. This demonstrated that of 201 derepressed genes in *Smchd1^MD1/MD1^* MEFs, 139 (69%) were repressed on the Xi in *Smchd1^MD1/MD1^* EpiSCs, suggesting that most of the X-linked genes derepressed in mutant MEFs had undergone inactivation in the undifferentiated epiblast even in the absence of SmcHD1 (Sakakibara *et al.*, 2018). It is worth mentioning, however, that genes depressed in *Smchd1^MD1/MD1^* MEFs tended to have relatively high Xi-probability, even though they were classified as inactivated genes (% Xi < 10%) in *Smchd1^MD1/MD1^* EpiSCs (Figure 2C), implying that they were barely repressed. In addition, these genes with relatively higher Xi-probability were not clustered but rather were distributed chromosome-wide (Figure 2D). These findings suggest that the repressive state of the Xi in *Smchd1^MD1/MD1^* EpiSCs was not as robust as that in wild-type EpiSCs.

**Figure 2.**
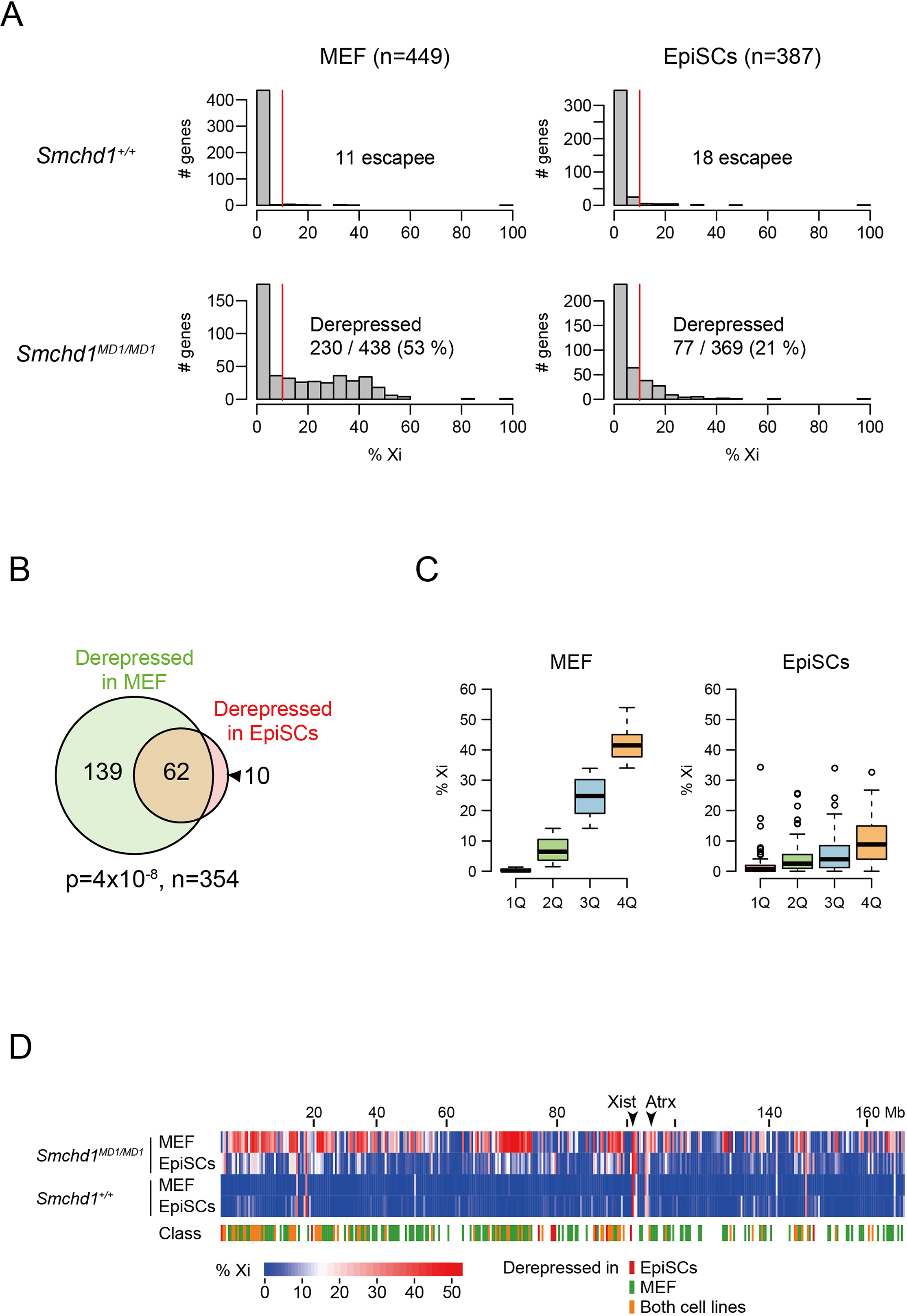
Chromosome-wide analysis of gene expression on the Xi in *Smchd1^MD1/MD1^* EpiSCs. (A) Histograms showing the numbers of genes with the respective scores of Xi-probability (% Xi) in wild-type (top) and *Smchd1^MD1/MD1^* (bottom) MEFs (left) and those EpiSCs (right) on *Xist^ΔA^*/+ background. A red line indicates a threshold (% Xi = 10%) whether genes were expressed on the Xi. (B) Venn diagram showing the number of genes derepressed in mutant MEFs and EpiSCs among the 354 informative genes commonly silenced in both wild-type EpiSC lines and their overlap. p-value, chi-squared test. (C) Three hundred and fifty-four informative genes commonly repressed on the Xi in wild-type MEFs and EpiSCs but derepressed in mutant MEFs were classified into four quartile groups according to the Xi probabililty (% Xi) in the mutant MEFs. Box plot shows the distribution of % Xi in each of the four quartile groups in mutant MEFs (left) and that in those groups in mutant EpiSCs (right). (D) Heatmap showing % Xi of respective informative X-linked genes along the chromosome in wild-type and *Smchd1^MD1/MD1^* MEFs and EpiSCs. Class indicates which category in the Venn diagram shown in (B) each X-linked gene belongs to in a color-matched way with the Venn diagram.

### H3K27me3 and H2AK119ub on the Xi visualized by immunostaining were more intensely stained in *Smchd1^MD1/MD1^* EpiSCs than in wild-type EpiSCs

The above findings prompted us to study the epigenetic state of the Xi in EpiSCs. It has been shown that the Xi in female mouse somatic cells is enriched for repressive histone modifications such as H3K27me3 and H2AK119ub, which can readily be detected by immunofluorescence using antibodies against these modifications (Fang *et al*, 2004; Plath *et al*, 2003; Silva *et al*, 2003; Erhardt *et al*, 2003; Napoles *et al*, 2004). Although another repressive histone modification, H3K9me3, is also enriched on the Xi in human and other mammals (Chadwick & Willard, 2004; Chaumeil *et al*, 2011; Rens *et al*, 2010; Zakharova *et al*, 2011; Nozawa *et al*, 2013), it is known to be undetectable in the mouse by immunofluorescence (Sado & Sakaguchi, 2013; Chaumeil *et al*, 2011), although more recent ChIP-seq has revealed enrichment of H3K9me3 on the mouse Xi (Keniry *et al*, 2016). We first performed immunostaining of wild-type and *Smchd1^MD1/MD1^* EpiSCs using antibodies against H3K27me3 and H2AK119ub. It was evident that both modifications formed one large overlapping accumulation in the nucleus in not only wild-type but also *Smchd1^MD1/MD1^* EpiSCs, suggesting that these modifications were enriched on the Xi in the mutant EpiSCs (Figure 3A). Since fluorescence for these modifications appeared rather intense in the mutant EpiSCs, we compared the signal intensity of these modifications between wild-type and mutant EpiSCs. The results demonstrated that immunofluorescence for H3K27me3 and H2AK119ub were both significantly higher in the mutant than in wild-type EpiSCs (Figure 3B). This was also the case in the comparison between *Smchd1^MD1/MD1^* and wild-type MEFs (Figure 3A and B). Western blotting, however, demonstrated that global levels of these modifications were comparable between *Smchd1^MD1/MD1^* and wild-type EpiSCs and also between the corresponding MEFs (Supplementary Figure S2).

**Figure 3.**
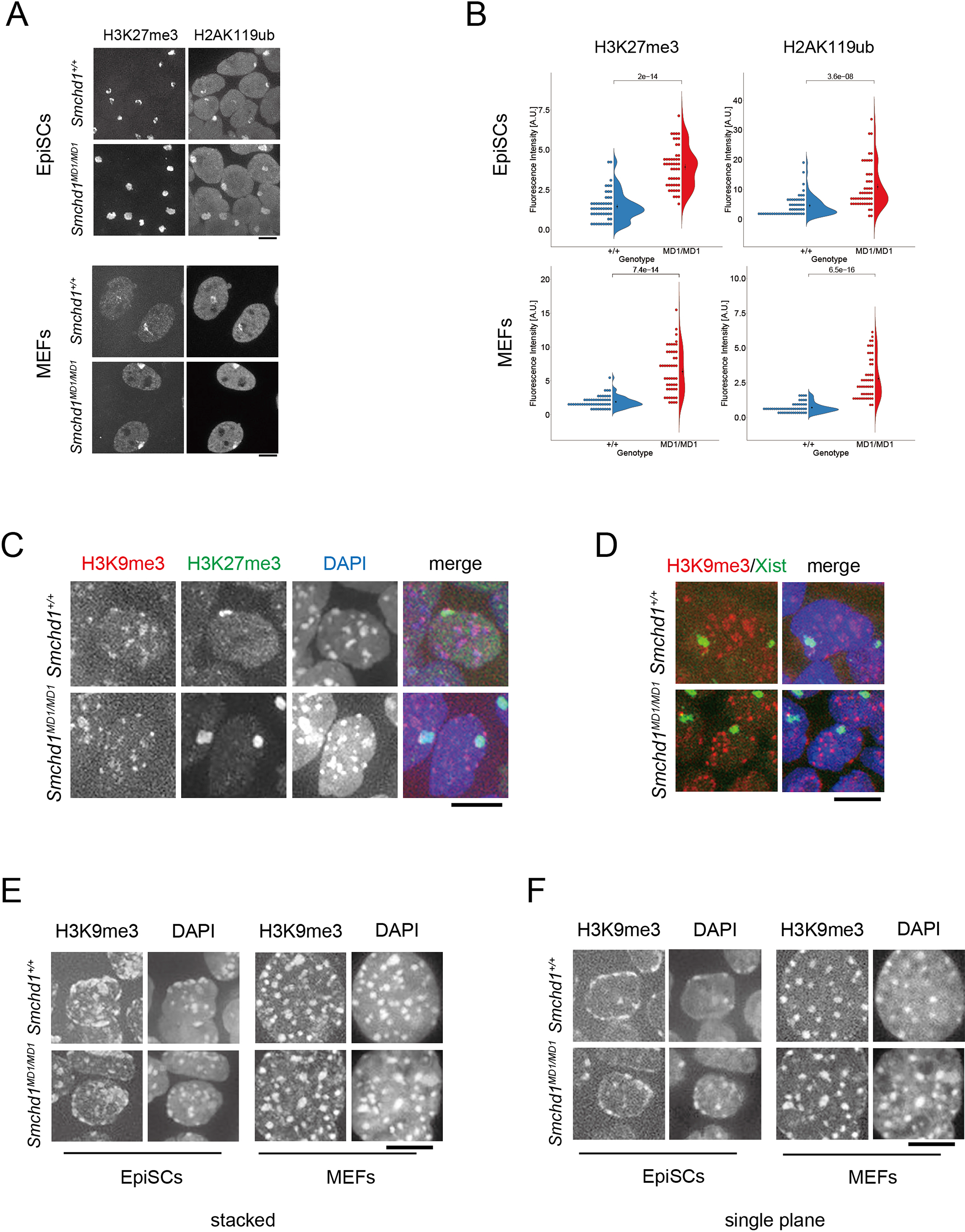
Histone modifications in EpiSCs and MEFs of wild-type and *Smchd1^MD1/MD1^.* (A) Immunofluorescence of wild-type and *Smchd1^MD1/MD1^* EpiSCs and MEFs for H3K27me3 and H2AK119ub. Both modifications were enriched on the inactive X chromosome regardless of the genotype. (B) Signal intensity of the immunofluorescence produced by the antibodies against H3K27me3, H2AK119ub, and H3K9me3 were compared between wild-type (blue) and *Smchd1^MD1/MD1^* (red) EpiSCs (top) and their MEF counterparts (bottom). p-value, Wilcoxon-Mann-Whitney test. (C) Immunofluorescence of wild-type and *Smchd1^MD1/MD1^* EpiSCs for H3K9me3 in combination with H3K27me3. (D) Immuno-RNA-FISH of wild-type and *Smchd1^MD1/MD1^* EpiSCs for H3K9me3 (red) and *Xist* RNA (green). (E) Immunofluorescence of wild-type and *Smchd1^MD1/MD1^* EpiSCs for H3K9me3 in comparison with that of wild-type and *Smchd1^MD1/MD1^* MEFs. Multiple images were stacked into one by projection. (F) Images of one single plane drawn from the respective stacked images shown in (E). Scale bar: 10 μm.

We also performed immunostaining for H3K9me3 in combination with H3K27me3. The Xi highlighted with antibody against H3K27me3 in both wild-type and *Smchd1^MD1/MD1^* EpiSCs was barely stained with an antibody against H3K9me3 (Figure 3C, Supplementary Figure S3). This was also the case when the Xi was visualized by RNA-FISH detecting *Xist* RNA in combination with immunostaining against H3K9me3 (Figure 3D). These results demonstrated that, as was the case in MEFs, H3K9me3 was not detectable by immunofluorescence on the Xi in EpiSCs. Although this was not necessarily unanticipated, it was of interest to find that accumulation of H3K9me3 in EpiSCs was distinct in shape and distribution in the nucleus from that in MEFs. While immunofluorescence signals produced by the antibody in MEFs were discrete and located more interiorly in the nucleus, forming several distinct chromocenters, they were often fused and were distributed at the periphery of the nucleus in EpiSCs (Figure 3E and 3F). In addition, although the staining of H3K9me3 basically colocalized with DAPI-dense heterochromatin, it often appeared to spread in a larger area than DAPI-dense heterochromatin in EpiSCs (Figure 3E and 3F). These results suggested that DAPI-dense heterochromatin in EpiSCs was different in nature from that in differentiated cells such as MEFs.

### Lrif1 does not accumulate on the Xi in EpiSCs and its association with DAPI-dense heterochromatin is lost in the absence of SmcHD1

It has been suggested that both SmcHD1 and its interacting factor Lrif1, a mouse ortholog of human HBiX1, accumulate on the Xi and facilitate chromatin condensation in differentiated female cells (Nozawa *et al*, 2013; Brideau *et al*, 2015). In female MEFs, antibodies raised against each of these proteins produced an intense signal on the Xi (Figure 4A, B, Supplementary Figure S4, and S5). Lrif1 also localized on DAPI-dense heterochromatin forming chromocenters, although to a lesser extent as compared to its localization on the Xi, whereas SmcHD1 was detected on chromocenters only in a minor fraction of cells (Supplementary Figure S4). We examined the cellular localization of SmcHD1 and Lrif1 in EpiSCs using the respective antibodies in combination with anti-H3K27me3. Immunofluorescence signals for SmcHD1 were rather intense and detected on not only the Xi marked with H3K27me3 but also DAPI-dense heterochromatin in wild-type EpiSCs (Figure 4A and Supplemental Figure S4). On the other hand, the signals for Lrif1 were detected at the DAPI-dense heterochromatin but largely off the Xi highlighted with H3K27me3 (Figure 4B and Supplementary Figure S5). Interestingly, Lrif1 lost its specific localization in the nuclei of both EpiSCs and MEFs homozygous for *Smchd1^MD1^* (Figure 4B). This was not due to downregulation of Lrif1, as similar levels of the protein were detected in wild-type and *Smchd1^MD1/MD1^* EpiSCs and MEFs by western blotting (Figure 4C). These results suggest that Lrif1’s localization to DAPI-dense heterochromatin as well as to the Xi depends on SmcHD1. The reason why Lrif1 does not appreciably localize to the Xi despite the presence of SmcHD1 on the Xi in wild-type EpiSCs is currently unknown. Nonetheless, given the differences in the localization of Lrif1 on the Xi as well as the spatial distribution of H3K9me3 in the nucleus between MEFs and EpiSCs in wild type, it seems that neither constitutive heterochromatin nor the facultative heterochromatin, that is, the Xi, are yet fully established in EpiSCs.

**Figure 4.**
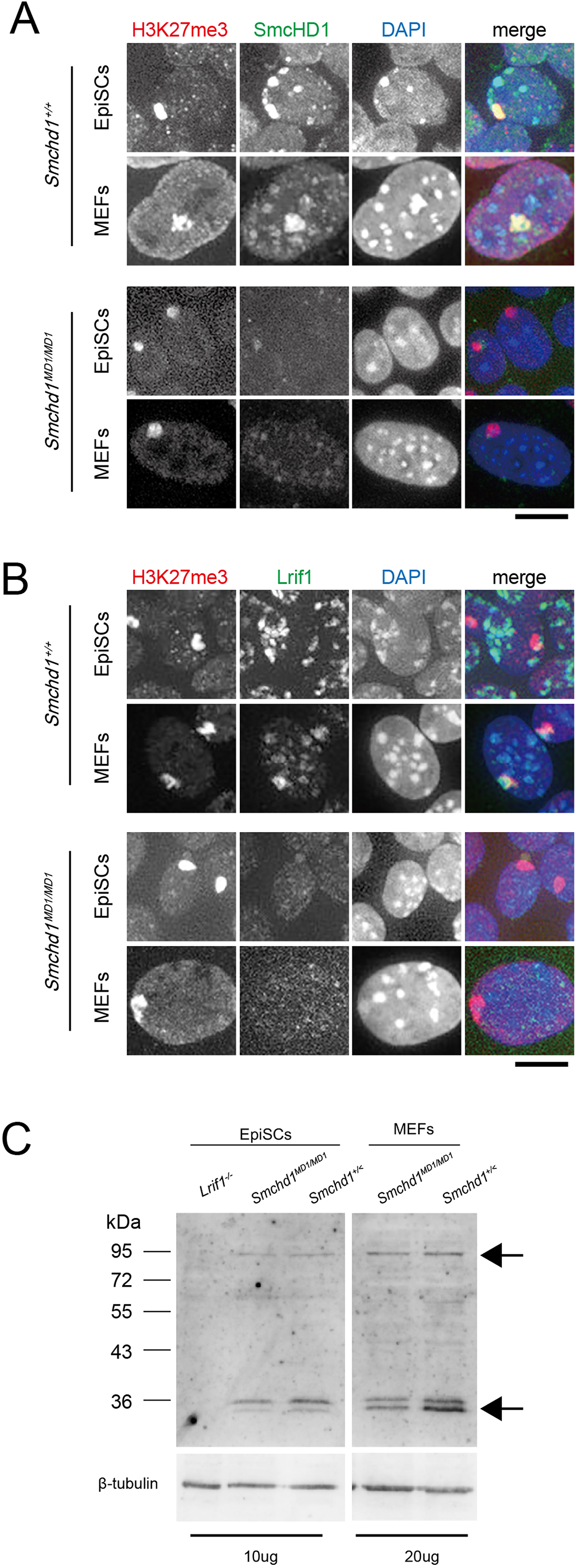
Cellular localization of SmcHD1 and Lrif1 in wild-type and *Smchd1^MD1/MD1^* EpiSCs and MEFs. (A) Immunofluorescence of wild-type and *Smchd1^MD1/MD1^* EpiSCs for SmcHD1 and H3K27me3 in comparison with that of wild-type and *Smchd1^MD1/MD1^* MEFs. (B) Immunofluorescence of wild-type and *Smchd1^MD1/MD1^* EpiSCs for Lrif1 and H3K27me3 in comparison with that of wild-type and *Smchd1^MD1/MD1^* MEFs. Scale bar: 10 μm. (C) Western blot showing Lrif1 expression levels in wild-type and *Smchd1^MD1/MD1^* EpiSCs (left) and those in wild-type and *Smchd1^MD1/MD1^* MEFs (right).

### SmcHD1-dependent X-linked genes with lower enrichment of H3K27me3 in mutant MEFs acquired H3K27me3 at the early stage of XCI

We previously demonstrated that X-linked genes that are derepressed in MEFs prepared from E13.5 *Smchd1^MD1/MD1^* fetuses showed lower enrichment of H3K27me3 and belong to a group of genes with relatively lower enrichment of H3K27me3 at the blastocyst stage of wild-type embryos (Sakakibara *et al*, 2018). This finding suggests that in *Smchd1^MD1/MD1^* embryos, these X-linked genes fail to acquire H3K27me3 or to maintain H3K27me3 acquired early on during postimplantation development due to the absence of SmcHD1 and eventually become derepressed. To explore these possibilities, we examined the H3K27me3 state in *Smchd1^MD1/MD1^* EpiSCs assuming their chromatin state to be comparable to that in the postimplantation epiblast lying midway between the blastocyst and mid-gestation stages. Chromatin immunoprecipitation (ChIP) using an antibody against H3K27me3, and subsequent quantitative PCR (qPCR) were carried out to see the enrichment of sequences at two X-linked loci, *Hprt* and *Rlim*, which were derepressed in *Smchd1^MD1/MD1^* MEFs but repressed in *Smchd1^MD1/MD1^* EpiSCs. The relative enrichment of H3K27me3 at these loci in *Smchd1^MD1/MD1^* EpiSCs was comparable to or even slightly higher than that in wildtype (Figure 5A). When the same ChIP-qPCR was performed using chromatin prepared from *Smchd1^MD1/MD1^* MEFs, lower enrichment of H3K27me3 was evident in *Smchd1^MD1/MD1^* MEFs than in wild type MEFs, being consistent with our previous ChIP-seq result reported by Sakakibara *et al* (2018). This finding suggests that these two X-linked genes acquire H3K27me3 at the early postimplantation development but fail to maintain it and become derepressed thereafter in the absence of SmcHD1.

**Figure 5.**
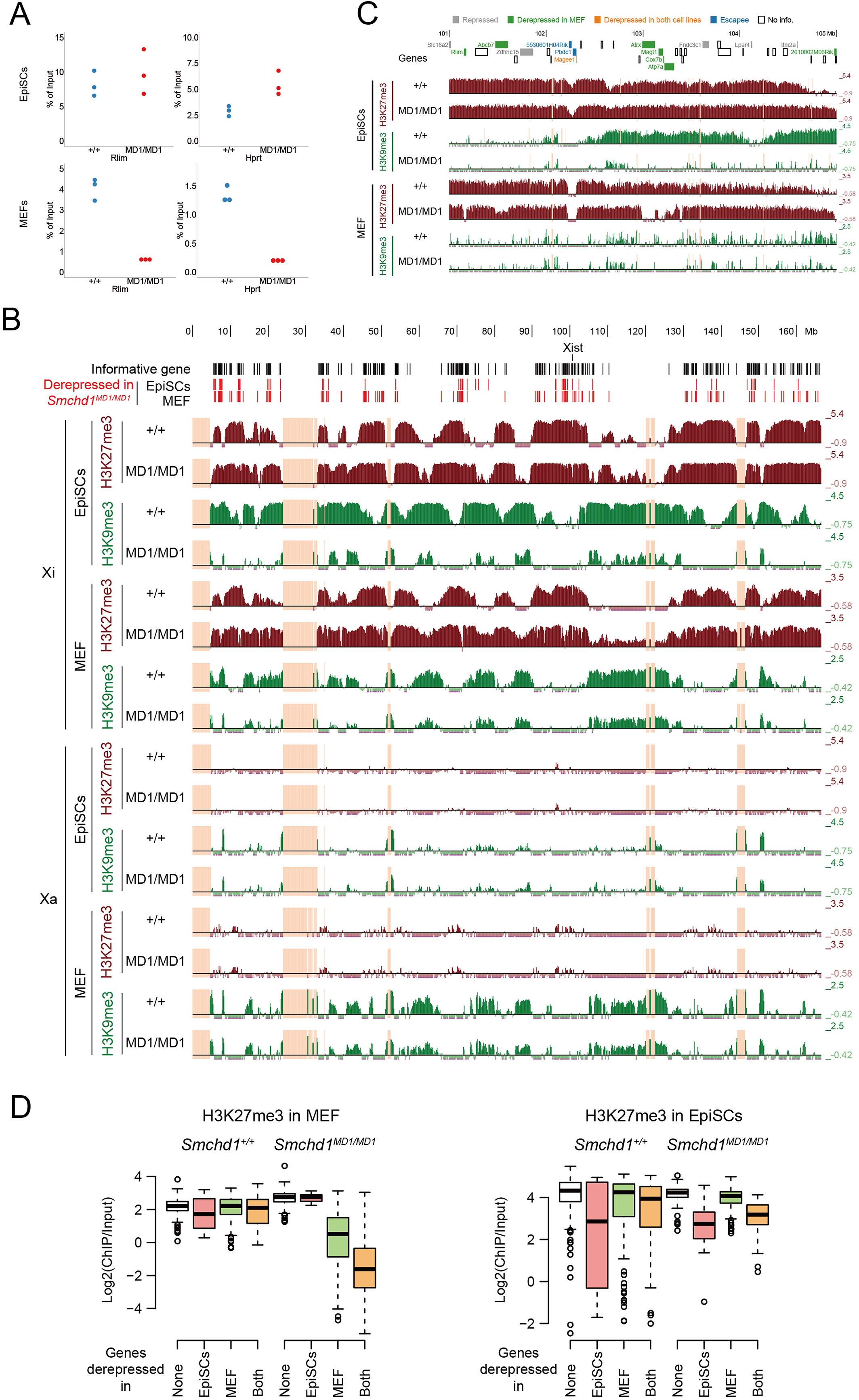
Histone modification states revealed by chromatin immunoprecipitation in wild-type and *Smchd1^MD1/MD1^ EpiSCs.* (A) ChIP against H3K27me3 and subsequent qPCR analysis at the *Rlim* and *Hprt* loci in wild-type (blue) and *Smchd1^MD1/MD1^* (red) EpiSCs in comparison with that in the respective MEFs. (B) Distribution of H3K9me3 and H3K27me3 on the Xi in wild-type and *Smchd1^MD1/MD1^* EpiSCs in comparison with that in the respective MEFs revealed by ChIP-seq. Enrichments of the respective histone modifications over the genome-wide average are shown as log2(ChIP/Input) per non-overlapping 150-kb bin in parallel with the locations of derepressed genes. Unmappable regions were indicated by light orange. (C) Distribution of H3K27me3 and H3K9me3 in the 4Mb region on the Xi containing *Atrx* in wild-type and *Smchd1^MD1/MD1^* EpiSCs and MEFs. Locations of genes in this region are shown on the top. The bin size was non-overlapping 5 kb. (D) Box plot showing H3K27me3 enrichment within gene bodies for each group of genes classified in Figure 2B according to the state of derepression (MEF-specific, EpiSC-specific, both, and not derepressed) in MEFs and EpiSCs. The color of each box matches the color of each group in the Venn diagram shown in Figure 2B and D.

### SmcHD1-deficiency results in substantial loss of H3K9me3 enrichment on the Xi in EpiSCs

To further study the impact of SmcHD1 deficiency on epigenetic regulation of the Xi in EpiSCs and MEFs, the chromosomal distribution of H3K27me3 and H3K9me3 was examined by allele-resolved ChIP-seq. H3K27me3 was almost absent on the Xa but was distributed along the Xi to form some blocks in both wild-type EpiSCs and MEFs, as expected (Figure 5B). On the other hand, while H3K9me3 was largely absent on the Xa, like H3K27me3, in EpiSCs, it was enriched at some limited regions on the Xa in MEFs. In addition, the Xi showed clear enrichment of H3K9me3 as some blocks in wild-type MEFs, in agreement with a previous report by Keniry *et al* (Keniry *et al*, 2016), and the same was true in wild-type EpiSCs (Figure 5B) even though immunostaining did not show obvious localization of H3K9me3 on the Xi in either case. These H3K9me3 blocks were distributed roughly in the regions with relatively lower enrichment of H3K27me3, although they partially overlapped with each other, on the Xi. In EpiSCs homozygous for *Smchd1^MD1^*, the distribution of H3K27me3 on the Xi was, at a glance, similar to that in wildtype, but, in fact, spread to some extent into the region normally occupied by H3K9me3 blocks (Figure 5B). The same trend was also seen in MEFs. Most interestingly, the enrichment of H3K9me3 was greatly diminished and became restricted only to limited regions on the Xi in the mutant EpiSCs. The situation was essentially the same in the mutant MEFs as well. In addition, the regions that still retained some enrichment of H3K9me3 on the Xi in the mutant EpiSCs coincided with those showing enrichment of H3K9me3 on the Xa (Figure 5B). It seems that these regions serve as hubs from which H3K9me3 spreads to form its blocks on the Xi, and this spreading is compromised in *Smchd1* mutant cells. The failure to form H3K9me3 blocks apparently allowed the expansion of H3K27me3 blocks into the adjacent regions normally occupied by H3K9me3. These results demonstrate that the most prominent effects caused by a functional loss of SmcHD1 in the mutant cells is substantial loss of H3K9me3 on the Xi.

### Effects of SmcHD1-deficiency on H3K27me3 on the Xi

To gain more insight into whether derepressed X-linked genes manifested by lower enrichment of H3K27me3 in homozygous mutant MEFs failed to acquire H3K27me3 from the beginning or lost it after acquiring it at the early developmental stages, the enrichment of H3K27me3 at the derepressed loci in mutant MEFs was compared between MEFs and EpiSCs. Figure 5C shows the distribution of H3K27me3 and H3K9me3 in a 4-Mb region on the Xi containing *Atrx* in EpiSCs in parallel with the distribution of these modifications in MEFs. *Atrx* and other genes such as *Rlim* in mutant EpiSCs, while they are derepressed with lower enrichment of H3K27me3 in the mutant MEFs, were silenced on the Xi (% Xi < 10%) with higher enrichment of H3K27me3. On the other hand, whether genes were embedded in H3K9me3-enriched regions or not in EpiSCs did not seem to be correlated with their ability to become derepressed in mutant MEFs. These observations suggest that the X-linked genes that initially undergo XCI but become derepressed in the absence of SmcHD1 acquire H3K27me3 at early developmental stages but lose it during later development. The box plot in Figure 5D shows that the genes that were derepressed only in mutant MEFs, the group of genes in green in Figure 2B and 5D, and those that stayed repressed in EpiSCs (the group of genes in white in Figure 5D) manifested comparably high enrichment of H3K27me3. They are, therefore, likely to be a class of genes that lose H3K27me3 acquired during the early phase of XCI at later developmental stages, suggesting that the Xi is defective in establishing the chromatin state that enables sustaining the H3K27me3 deposited at a subset of X-linked genes. On the other hand, there seemed to be no correlation between an enrichment of H3K9me3 and a class of genes derepressed in MEFs (Supplementary Figure S6). Given that SmcHD1-deficiency compromises proper distribution of H3K9me3 on the Xi, it is tempting to speculate that the lack of H3K9me3 might secondarily compromise proper accumulation and maintenance of H3K27me3 and eventually cause derepression of X-linked genes that were inactivated early on.

## Discussion

Several lines of evidence suggest that in female mouse embryos, SmcHD1 plays an important role in stable maintenance of the X-inactivated state during embryonic development. In the absence of SmcHD1, female embryos die during mid-gestation stages, at least partly due to placental defects caused by derepression of X-linked genes that had been inactivated early on. Since male embryos deficient for SmcHD1 are also perinatal lethal on the C57BL/6 background (Leong *et al*, 2013a), however, it is likely that SmcHD1 is critical for stable gene silencing in both sexes and females use this system for the maintenance of XCI as well. Although our previous study suggested that SmcHD1 is involved in establishment of epigenetic states required for stable silencing of X-inactivated genes, it has remained unaddressed how the lack of SmcHD1 affects the dynamics of histone modifications during development. To understand the details of the mechanism by which SmcHD1 contributes to establishing the epigenetic state of the Xi, we took advantage of EpiSCs derived from the undifferentiated postimplantation epiblast, in which XCI has already been initiated but is thought to be midway to the fully established state seen in MEFs. The idea that chromatin in EpiSCs is in a transitional state or premature as compared to that in MEFs was supported by the observations that Lrif1, known to colocalize with SmcHD1 on the Xi in MEFs, has not yet been localized on the Xi in EpiSCs and that DAPI-dense heterochromatin in EpiSCs tends to distribute at the periphery of the nucleus, contrasting with that forming discrete chromocenters more interiorly in the nucleus of MEFs and other differentiated cells in mice.

Allele-resolved RNA-seq of wild-type and *Smchd1^MD1/MD1^* EpiSC revealed that the majority of informative genes on the Xi were classified as being repressed and only about 20% of them were misexpressed on the Xi in *Smchd1^MD1/MD1^* EpiSC. This contrasts with the fact that more than half of the informative genes on the Xi were derepressed in *Smchd1^MD1/MD1^* MEFs. Although it is not clear whether those genes misexpressed in the mutant EpiSCs failed to be inactivated from the beginning or were once inactivated but have become derepressed, it is clear that genes on the Xi have been substantially silenced in *Smchd1^MD1/MD1^* EpiSCs. In addition, while these silenced genes manifested enrichment of H3K27me3, a subset of them were derepressed and manifested lower enrichment of H3K27me3 than the genome-wide average in *Smchd1^MD1/MD1^* MEFs. These findings suggest that during development of *Smchd1^MD1/MD1^* embryos, genes on the X chromosome coated with *Xist* RNA undergo inactivation and acquire H3K27me3 in the epiblast cells even in the absence of SmcHD1, but subsequently lose H3K27me3 and become derepressed, although it is not clear whether the loss of H3K27me3 precedes derepression or the other way around. In any case, it seems important to sustain H3K27me3 deposited at the silenced gene loci for the stable maintenance of the X-inactivated state.

An obvious anomaly found in the chromatin of the Xi in *Smchd1^MD1/MD1^* EpiSCs in comparison with wild type was extensive loss of H3K9me3. It should be noted, however, that H3K9me3 still showed some enrichment at limited regions on the Xi, which apparently coincided with the regions enriched in H3K9me3 on the Xa. It is, therefore, possible that SmcHD1-deficiency in EpiSCs does not necessarily compromise the deposition of H3K9me3 on the Xi per se but the spreading of its wave from such initially deposited sites or hubs. Nonetheless, given that the majority of genes on the Xi are silenced in *Smchd1^MD1/MD1^* EpiSCs, resembling their status in the postimplantation epiblast that has just undergone XCI, it is likely that H3K9me3 is not involved in the initial silencing of the genes by XCI. Several lines of evidence suggest that *Xist*-mediated silencing is triggered by deacetylation of H3K27ac at enhancers and followed by deposition of PRC1-mediated H2AK119ub and subsequently by PRC2-mediated deposition of H3K27me3 on the Xi (Żylicz *et al*, 2019; Nesterova *et al*, 2019). It has been shown that SmcHD1 is recruited to the Xi by H2AK119ub (Jansz *et al*, 2018b). In addition, localization of SmcHD1 on the Xi is apparently preceded by accumulation of H2AK119ub and H3K27me3 during differentiation of ESCs (Gendrel *et al*, 2014; Sakata *et al*, 2017). Taking these previous findings all together, we suggest that SmcHD1 plays a critical role in the propagation of H3K9me3 from the hubs on the Xi that has already been enriched with H2AK119ub and H3K27me3, and facilitates the establishment of the respective heterochromatin blocks of H3K27me3 and H3K9me3 on the Xi in the epiblast cells prior to differentiation (Figure 6). It seems that forming H3K9me3 blocks on the Xi restricts the occupancy of H3K27me3, and its failure disturbs the overall distribution of H3K27me3, resulting in irregular expansion of the H3K27me3 blocks, as seen in *Smchd1^MD1/MD1^* EpiSCs. If the amount of PRC2 available for the maintenance of H3K27me3 is limited, a subset of H3K27me3 deposited at the silenced X-linked gene loci early on might be partly lost, causing derepression of those loci over time. Alternatively, although this may not be mutually exclusive, propagation of H3K9me3 and subsequently formed blocks of H3K9me3 may facilitate further condensation of the Xi and render it very robust heterochromatin. In the absence of H3K9me3, the Xi could stay relatively less condensed, allowing access to demethylases of H3K27me3 at particular loci that have been silenced and their eventual derepression. Signal intensity of the Xi produced by immunofluorescence of antibodies against H3K27me3 and H2AK119ub was significantly enhanced in EpiSCs and MEFs homozygous for *Smchd1^MD1^* compared to their wild-type counterparts. This could be explained by apparent expansion of H3K27me3 blocks on the Xi in the mutant. It is also possible that the less-condensed chromatin state of the Xi in the mutant might give the respective antibodies easier access to their target, contributing to an increase in signal intensity. The proposed role of SmcHD1 in the higher order chromatin structure might thus be mediated by appropriate formation of H3K27me3 and H3K9me3 blocks. We previously showed that depletion of SmcHD1 by CRIPSPR/Cas9-mediated genome editing does not compromise the silencing state of the Xi in MEFs, suggesting that SmcHD1 is not essential for the maintenance of the X-inactivated state once it is established. In this case, both H3K27me3 and H3K9me3 blocks should have been formed on the Xi in the original MEFs by the time SmcHD1 was depleted. Formation of such H3K27me3 and H3K9me3 blocks should allow establishment of the epigenetic state and higher order chromatin structure required for stable maintenance of the X-inactivated state. It is likely that SmcHD1 plays an important role in establishing this epigenetic state and chromatin structure and they are maintained independently of SmcHD1 once established.

**Figure 6.**
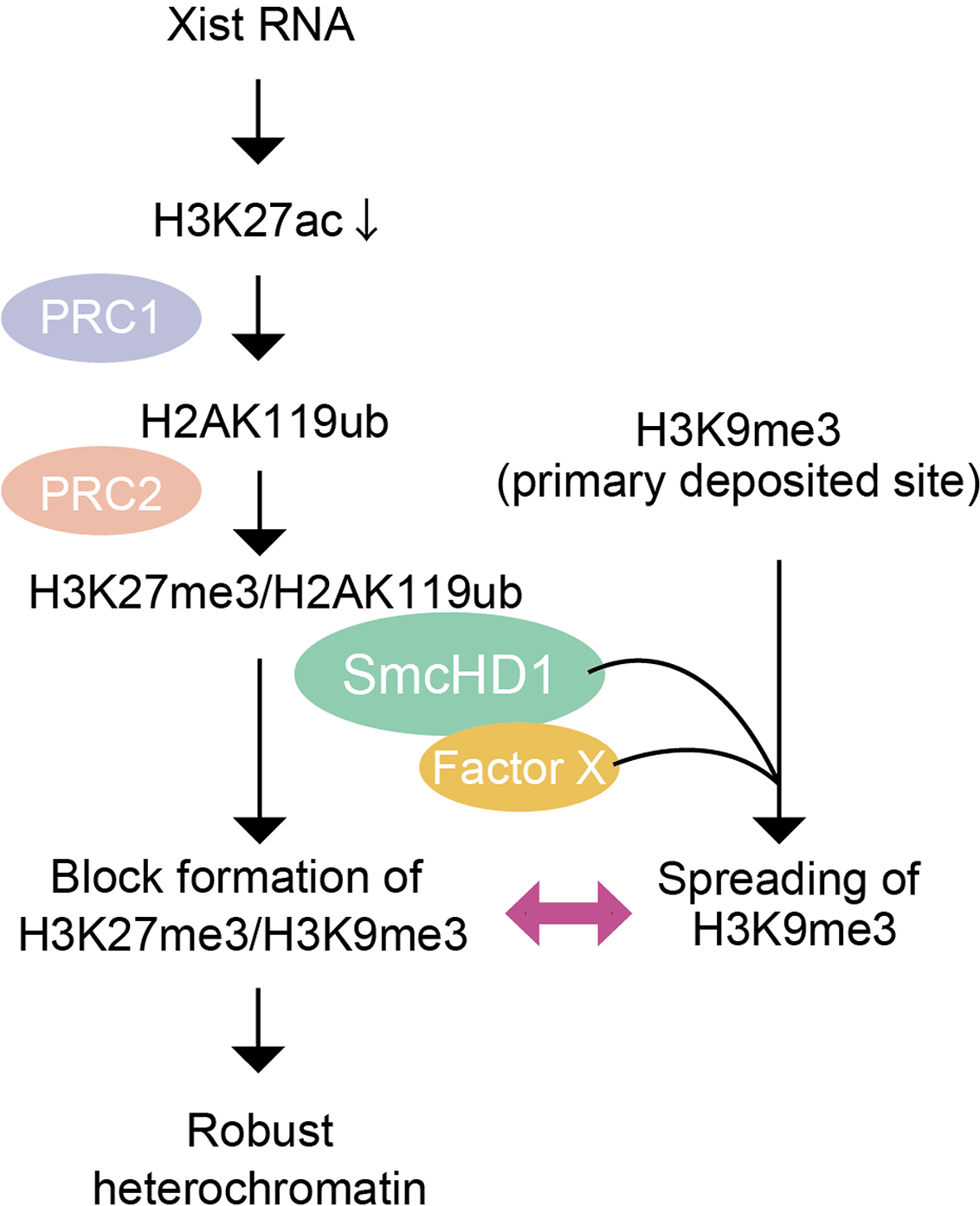
Model for how Xi acquires H3K27me3/H3K9me3 blocks in a SmcHD1-dependent manner. (A) The finding of this study was incorporated into the current view of how XCI is initiated and epigenetic modifications of the Xi are regulated.

It has been shown that H3K9me3 plays an important role in silencing of endogenous retroviruses (ERVs) and its loss caused by the lack of histone methyltransferase SetDB1 results in derepression of not only many classes of ERVs in mouse ES cells but also certain classes of ERVs even in differentiated cells (Kato *et al*, 2018; Matsui *et al*, 2010). Since the lack of SmcHD1 caused extensive loss of H3K9me3 on the Xi, we wondered if silencing of ERVs was affected in *Smchd1^MD1/MD1^* EpiSCs. We did not, however, find evidence that the lack of SmcHD1 caused derepression of transposons in mutant EpiSCs (data not shown).

As far as we are aware, there has been no direct evidence showing an interaction between SmcHD1 and machinery regulating H3K9me3. A possible link between SmcHD1 and H3K9me3, however, might be envisioned based on previous findings, including ours, that SmcHD1 could associate with HP1, a protein that recognizes and binds H3K9me3, via an interaction with Lrif1 (or HBiX1 in human) (Nozawa *et al*, 2013; Brideau *et al*, 2015). SmcHD1 and Lrif1 localize at DAPI-dense heterochromatin enriched with H3K9me3 in EpiSCs. Interestingly, Lrif1 lost its association with DAPI-dense heterochromatin in the absence of SmcHD1 in both EpiSCs and MEFs, suggesting that Lrif1’s localization at DAPI-dense heterochromatin depended on SmcHD1. It is therefore reasonable to assume that SmcHD1 may have an alternative way to associate with DAPI-dense heterochromatin enriched for H3K9me3 without the interaction with HP1 via Lrif1. Candidates for the factors that associate with SmcHD1 and regulate H3K9me3 on the Xi may be found among factors known to participate in the regulation of H3K9me3 at DAPI-dense heterochromatin.

### Materials and methods Mice

Details of MommeD1 mice were described elsewhere (Blewitt *et al*, 2008) and the genetic background of the MommeD1 colony has been replaced essentially by C57Bl/6 (B6) (Leong *et al*, 2013b). Two types of *Xist* mutants carrying either *Xist^ΔA^* or *Xist^1lox^* were described previously (Sado *et al*, 2005; Hoki *et al*, 2009) and their genetic background has also been replaced essentially by B6. All of these mice were maintained by crossing heterozygous females with B6 males. Male mice heterozygous for *Smchd1^MD1^* with the X chromosome derived from the JF1 strain (X^JF1^) [X^JF1^Y; *Smchd1^+/MD1^*] were generated by crossing JF1 females with *Smchd1^+/MD1^* males. Females doubly heterozygous for *Smchd1^MD1^* as well as one of either *Xist^ΔA^* or *Xist^1lox^* were generated by crosses between females with one of the *Xist* mutations and males heterozygous for *Smchd1^MD1^*.These males and females were crossed to obtain [XB6-ΔAX^JF1^; *Smchd1^MD1/MD1^*] or [XB6-1loxX^JF1^; *Smchd1^MD1/MD1^*] embryos for derivation of EpiSCs and for preparation of MEFs. Primer sequences used for genotyping are shown in Supplementary Table S1. All mice were maintained and used in accordance with the Guidelines for the Care and Use of Laboratory Animals of Kindai University (KDAS-26-006).

### Cell culture

EpiSCs were established from E6.5 epiblasts in the presence of IWP2 according to the previous report (Sugimoto *et al*, 2015). For passage, cells were treated with 10 μM Y27632 for 1 hour prior to dissociation with Accutase. Dissociated cells were seeded on a layer of feeder cells in the presence of 10 μM Y27632, and the medium was exchanged daily. MEFs were prepared from fetuses obtained at E12.5 as described previously (Sakata *et al*, 2017).

### RNA-FISH

Plasmid pXist_SS12.9 containing a 12.9-kb *Xist* cDNA fragment (Sakata *et al*, 2017) and BAC clone RP23-260I15 containing the *Atrx* gene were labeled with Green dUTP and Cy3 dUTP, respectively, by nick translation. EpiSCs or MEFs grown on a coverslip were fixed with 4% paraformaldehyde (PFA) for 10 minutes. Following permeabilization, blocking in 0.05% Triton-X100/0.05% BSA/PBS for 30 minutes, and subsequent dehydration in 70% and 100% EtOH, air-dried specimens were subjected to hybridization as described by Sakata *et al* (Sakata *et al*, 2017). Cytological preparations using Carnoy’s fixative were also made and subjected to hybridization.

### Allelic expression analysis by RT-PCR

Total RNA was converted into cDNA using SuperScript III (Invitrogen) using random hexamer as a primer. RT-PCR was subsequently carried out to amplify the fragments containing restriction site polymorphisms using primer sets shown in Supplementary Table S1. PCR products were digested with an appropriate restriction enzyme (*Pdha1*, *Taq*I, *G6pd*, *Dra*I, *Hprt*, *Hin*fI, *Rex3*, *Bsr*GI, *Rbm3*, or *Dde*I).

### Immuno-RNA-FISH

Cells grown on a coverslip were fixed with 2% PFA/0.5% TritonX-100 for 10 min and subjected to permeabilization and blocking in 0.5% Triton-X100/0.5% BSA/PBS for 30 min. Immunoreaction was carried out using an antibody against H3K9me3 in the presence of RNase Out (Invitrogen) at 1 U/μL at 4°C overnight, and subsequently the primary antibody was visualized with CF594-labeled goat anti-mouse IgG antibody (Biotium). Following postfixation with 4% PFA, RNA-FISH was carried out as described above, with the dehydration process being skipped.

### Immunofluorescence

Cells were grown on a coverslip. Fixation and subsequent permeabilization and blocking were carried out in a slightly different way depending on the antibody used, as shown in Supplementary Table S2. Fluorescence images were taken with an inverted microscope (IX71, Olympus) equipped with an Olympus Disk Scanning Unit (DSU) and an EM-CCD camera (iXon, Andor), and analyzed with MetaMorph imaging software (Molecular Devices).

Antibody against Lrif1 was raised by immunizing rabbits with a His-tagged recombinant protein containing the last 237 amino acids of Lrif1 and affinity-purified using the antigen. The specificity of the antibody was confirmed using Lrif1. Antibodies used for this study are listed in Supplementary Table S2.

### Quantitation of fluorescence intensity

Immunofluorescence images were analyzed using ImageJ (1.52h) for comparison of signal intensity. Fluorescence of the Xi visualized as a large domain in the nucleus by using antibodies against either H3K27me3 or H2AK119ub was quantified and normalized as follows: (area of a selected domain produced by either antibody x mean fluorescence intensity of selected domain) - (area of the selected domain x mean fluorescence intensity of the nucleoplasm surrounding the selected domain). Wilcoxon-Mann-Whitney test and data visualization were conducted using R (4.0.1)

### Allele-resolved RNA-seq

EpiSCs grown on feeder cells were treated with Y27632 and dissociated into single cells using Accutase. The cell suspension was transferred onto a gelatin-coated culture dish and incubated for 30 min at 37 °C to remove feeder cells based on their differential adherence. Following one more round of removal of the feeder cells, EpiSCs were collected by centrifugation and disrupted in TRIzol (Ambion). Total RNA was prepared according to the manufacturer’s instructions. Libraries were prepared using the TruSeq Stranded mRNA LT Sample Prep Kit (Illumina). Paired-end 101 bp sequences were obtained using the Illumina platform.

Allele-resolved RNA-seq analysis was performed as described by Sakata *et al.* (2017) with minor modifications: (1) paired-end reads were used as single-end reads; (2) the numbers of allele-specific reads in replicates were summed; (3) genes with more than 10 allele-specific reads were considered to be informative for assessing the expression from the Xi; (4) Cufflinks v2.2.1 cuffdiff (Trapnell *et al*, 2010) with option ‘--library-type ff-firststrand --multi-read-correct’ was used to estimate gene expression level as the FPKM (fragments per kilobase of exon per million fragments mapped) in a non-allelic manner. RNA-seq data of MEFs were reported previously (Sakakibara *et al*, 2018). For each gene, the number of maternal (B6 or 129) and paternal (JF1) allele-specific reads were counted and percentages of total expression from the paternal allele were calculated as %Xi for assessing the expression from the Xi. According to a generally accepted threshold (Peeters *et al.*, 2014), genes with ≥ 10% of total expression from the Xi in wild type were classified as escapees. To distinguish repressed genes from de-repressed genes in non-escapees, we set 10% of total expression from the Xi as a threshold. For comparison among cell lines, only genes with informative % Xi in all cell lines to be compared were considered. For Figure S1D, genes with FPKM ≥ 1 were considered to be expressed.

### Western blotting

EpiSCs free from feeder cells were suspended in RIPA buffer/1x protease inhibitor cocktail and placed on ice for 10 minutes. Following centrifugation at 15,000 rpm for 10 minutes at 4°C, the supernatant was collected. SDS-PAGE and immunoreactions were carried out using standard protocols. Images were taken with an ImageQuant LAS 500 (GE Healthcare Life Sciences).

### Chromatin immunoprecipitation and allele-resolved ChIP-seq

For ChIP-qPCR, MEFs and EpiSCs were fixed with DMEM containing 5% formaldehyde. Preparation of chromatin and immunoprecipitation using an antibody against H3K27me3 were carried out as previously described (Nozawa *et al*, 2013). The primers used for ChIP-qPCR are shown in Supplementary Table S1.

For ChIP-seq, following chromatin immunoprecipitation using antibodies against either H3K27me3 or H3K9me3, libraries were generated using the NEBNext Ultra II DNA Library Prep Kit (New England Biolabs) according to the manufacturer’s instructions. Size fractionation was carried out to enrich fragments of 100-300 bp from purified DNA for library preparation as well as the library around the size of 200 bp fragments after PCR amplification using AMPure XP (Beckman Coulter) magnetic beads. Each library thus size-fractionated was quantified using a LabChip GX (Perkin Elmer). Libraries were pooled and sequenced on the Illumina HiSeq 3000 to produce 100 bp paired-end reads.

#### Analysis of allele-resolved ChIP-seq datasets

Paired-end reads were trimmed to remove adaptor sequences using the cutadapt (Martin 2011) and FLASH2 (Magoc and Salzberg, 2011) before mapping, and those with ≥ 50 bp were used as single-end reads for further analysis. The trimmed reads were mapped to B6^Xist_mutant^/JF1 strain-specific diploid genome as described by Sakata *et al.* (Sakata *et al*, 2017) using bwa samse (Li & Durbin, 2009). The custom reference genome included the chromosomes of JF1, the X chromosome of 129S1, and the B6 mm9 reference genome. The mapped positions were converted to mm9 genomic coordinates keeping the strain origin because our custom reference genome incorporated indels in addition to SNPs. The mapping quality was recalculated to be equivalent to the haploid reference by counting the number of suboptimal hits per haploid genome, as the diploid genome has twice as many suboptimal alignments as those on the haploid genome (Miura *et al*, 2020). Reads with low mapping quality (<20) were discarded to ensure that reads mapped to unique genomic positions were considered with confidence. Possible PCR duplicates were removed using samtools (Li *et al*, 2009). As a result, each mapped read had a single coordinate of the mm9 reference and a set of possible strain origins.

Note that, for the cell lines used in this study, the genomic sequence of the 129 strain was retained at the vicinity of the mutated *Xist* locus even after extensive backcrosses into the B6 background, and not all autosomal regions were B6/JF1 hybrid due to the derivation of the F2 generation (see Fig. 1A). Therefore, the haplotype of each cell line was required to assign the parental origin of a read using a set of possible strain origins at each SNP/indel site. The haplotypes of chromosome X were inferred as described previously (Sakata *et al*, 2017). For the autosomal haplotypes, regions with B6/JF1 hybrid were roughly determined by the ratio of JF1-allele to total allele-specific read counts for each 500 kb bin in the input sample, and the boundaries between hybrid and non-hybrid regions were inferred by visual inspection of the distribution of JF1-specific reads in the input and the ChIP samples. Using the haplotype for each cell line, we counted maternal-specific, paternal-specific, and all mapped reads separately in the ChIP and input samples in non-overlapping bins across the chromosome and expressed them as RPKM (reads per kilobase per million mapped reads). We calculated the enrichment ratio of the ChIP sample to the corresponding input defined as log2[(ChIP RPKM)/(input RPKM)]. Bins with RPKM < 0.1 for all mapped reads and RPKM < 0.02 for parental-specific reads were excluded as unmappable. To facilitate direct comparison of the distribution of the histone modification between the cell lines, each profile of the ChIP enrichment was normalized to that of wild-type MEF so that the mode of the difference between the two autosomal profiles was 0. For each histone modification in each cell line, a single normalization factor was determined using the non-allelic profile of autosomal 50 kb bins and applied to any profiles regardless of chromosome, bin size, or parental origin. The mode was estimated by the R package ‘modeest’.

We used the UCSC genome browser to visualize the ChIP enrichments on genomic coordinates (Kent *et al*, 2002) with scales indicated in each track and the window option ‘mean’. To avoid interpreting unmappable bins as unenriched, those ChIP enrichments were estimated by linear interpolation in the case of a single unmappable bin flanked by mappable bins. Otherwise, in the case of successive unmappable bins, we filled those bins with light orange in all figures. To analyze the ChIP enrichment for each X-linked gene, we defined its gene body as a bin and excluded its length < 1 kb due to low read counts.

## Data availability

The allele-specific RNA-seq and ChIP-seq data generated in this study have been deposited in the NCBI Gene Expression Omnibus (GEO) database under accession code XXXXXX.

## Acknowledgments

This work was supported partly by Grants-in-Aid for Scientific Research (A) from the Japan Society for the Promotion of Science (JSPS) to TS (17H01588 and 20H00550) and C.O. (19H03156), respectively, Grants-in-Aid for Scientific Research on Innovative Areas from the Ministry of Education, Culture, Sports, Science and Technology (MEXT) (17H06426 to K.N; 18H05532 to C.O.), and a Takeda Science Foundation grant to T.S. We would also like to thank Tetsuya Hori and Shiho Ogawa for technical advice about ChIP-seq library preparation and Rawin Poonperm for imaging analysis.

---

**Supplementary Figure 1. Effect of SmcHD1-deficiency on X-inactivated genes was essentially the same between Xist^ΔA^/+ and Xist^1lox^/+ background in EpiSCs.**

(A) Expression levels of marker genes for EpiSCs, ESCs, and MEFs were compared in EpiSCs and MEFs with respective genotypes. Expression profile of these marker genes in ESCs was extracted from our recent publication (Takahashi et al, 2019).

(B) Histograms showing the numbers of genes with the respective scores of Xi-probability in wild-type (top) and *Smchd1^MD1/MD1^* (bottom) EpiSCs on *Xist^1lox^*/+ background.

(C) Comparison of the degree of derepression for X-linked genes on the Xi between *Xist^ΔA^*/+ and *Xist^1lox^*/+ background. (left) Of 377 informative genes in all 4 relevant EpiSC lines, 344 genes (black open circles) were silenced on the Xi and 12 genes (blue filled circles) were escaped from XCI in both wild-type EpiSC lines. Sixteen (red open circles) and 5 (green open circles) genes were classified as escapees in wild type on *Xist^1lox^*/+ and *Xist^ΔA^*/+ background, respectively. (right) Although the number of escaped genes in wild type was larger on *Xist^1lox^*/+ background than on *Xist^ΔA^*/+, 344 commonly silenced genes were similarly derepressed in *Smchd1^MD1/MD1^* on both *Xist* background (Pearson correlation coefficient, r = 0.86).

(D) Cumulative distribution plot of fold-changes of expressed X-linked genes (red) versus autosomal genes (black) between wild-type and *Smchd1^MD1/MD1^* in MEFs and respective EpiSCs lines as indicated.

**Supplementary Figure 2. Global levels of histone modifications in EpiSCs and MEFs.**

Western blot analysis showing global levels of H3K27me3 and H2AK119ub in wild-type and *Smchd1^MD1/MD1^* EpiSCs and their respective counterparts in MEFs. Twenty micrograms of whole cell lysate was loaded.

**Supplementary Figure 3. Immunofluorescence of H3K9me3 and H3K27me3 in wild-type and Smchd1^MD1/MD1^ EpiSCs.**

A wider view of the field surrounding the nucleus shown in Figure 3C. The nucleus shown in Figure 3C is indicated by a rectangle in the merged image. Scale bar: 10 μm.

**Supplementary Figure 4. Immunofluorescence of SmcHD1 in wild-type EpiSCs and MEFs.**

A wider view of the field surrounding the nucleus shown in Figure 4A. The nucleus shown in Figure 4A is indicated by a rectangle in the merged image. While SmcHD1 localized to both the Xi marked with H3K27me3 and DAPI-dense heterochromatin in EpiSCs, its localization in MEFs was essentially to the Xi in most nuclei, and in a small fraction of nuclei, it localized to the DAPI-dense heterochromatin. Scale bar: 10 μm.

**Supplementary Figure 5. Immunofluorescence of Lrif1 in wild-type and *Smchd1^MD1/MD1^* EpiSCs and MEFs.**

A wider view of the field surrounding the nucleus shown in Figure 4B. The nucleus shown in Figure 4B is indicated by a rectangle in the merged image. While Lrif localized to both the Xi marked with H3K27me3 and DAPI-dense heterochromatin in both EpiSCs and MEFs, its localization was substantially lost in both EpiSCs and MEFs in the absence of SmcHD1. Scale bar: 10 μm.

**Supplementary Figure 6. The enrichment of H3K9me3 showed no correlation with derepression of X-linked genes.**

Box plot showing H3K9me3 enrichment within gene bodies for each group of genes classified in Figure 2B according to the state of derepression (MEF-specific, EpiSC-specific, both, and not derepressed) in MEFs and EpiSCs. The color of each box matches the color of each group in the Venn diagram shown in Figure 2B and D.

## References

Blewitt ME, Gendrel A-V, Pang Z, Sparrow DB, Whitelaw N, Craig JM, Apedaile A, Hilton DJ, Dunwoodie SL, Brockdorff N, et al (2008) SmcHD1, containing a structural-maintenance-of-chromosomes hinge domain, has a critical role in X inactivation. Nature Genetics 40

Borsani G, Tonlorenzi R, Simmler MC, Dandolo L, Arnaud D, Capra V, Grompe M, Pizzuti A, Muzny D, Lawrence C, et al (1991) Characterization of a murine gene expressed from the inactive X chromosome. Nature 351: 325–329

Brideau NJ, Coker H, Gendrel A-VV, Siebert CA, Bezstarosti K, Demmers J, Poot RA, Nesterova TB & Brockdorff N (2015) Independent Mechanisms Target SMCHD1 to Trimethylated Histone H3 Lysine 9-Modified Chromatin and the Inactive X Chromosome. Molecular and cellular biology 35: 4053–68

Brockdorff N, Ashworth A, Kay GF, Cooper P, Smith S, McCabe VM, Norris DP, Penny GD, Patel D & Rastan S (1991) Conservation of position and exclusive expression of mouse Xist from the inactive X chromosome. Nature 351: 351329a0

Brons GI, Smithers LE, Trotter MW, Rugg-Gunn P, Sun B, Lopes SM de, Howlett SK, Clarkson A, Ahrlund-Richter L, Pedersen RA, et al (2007) Derivation of pluripotent epiblast stem cells from mammalian embryos. Nature 448: 191–195

Chadwick BP & Willard HF (2004) Multiple spatially distinct types of facultative heterochromatin on the human inactive X chromosome. P Natl Acad Sci Usa 101: 17450–17455

Chaumeil J, Waters PD, Koina E, Gilbert C, Robinson TJ & Graves JAM (2011) Evolution from XIST-Independent to XIST-Controlled X-Chromosome Inactivation: Epigenetic Modifications in Distantly Related Mammals. Plos One 6: e19040

Chen G, Schell JP, Benitez JA, Petropoulos S, Yilmaz M, Reinius B, Alekseenko Z, Shi L, Hedlund E, Lanner F, et al (2016) Single-cell analyses of X Chromosome inactivation dynamics and pluripotency during differentiation. Genome Res 26: 1342–1354

Csankovszki G, Nagy A & Jaenisch R (2001) Synergism of Xist Rna, DNA Methylation, and Histone Hypoacetylation in Maintaining X Chromosome Inactivation. J Cell Biology 153: 773–784

Erhardt S, Su I, Schneider R, Barton S, Bannister AJ, Perez-Burgos L, Jenuwein T, Kouzarides T, Tarakhovsky A & Surani MA (2003) Consequences of the depletion of zygotic and embryonic enhancer of zeste 2 during preimplantation mouse development. Development 130: 4235–4248

Fang J, Chen T, Chadwick B, Li E & Zhang Y (2004) Ring1b-mediated H2A Ubiquitination Associates with Inactive X Chromosomes and Is Involved in Initiation of X Inactivation. J Biol Chem 279: 52812–52815

Gdula MR, Nesterova TB, Pintacuda G, Godwin J, Zhan Y, Ozadam H, McClellan M, Moralli D, Krueger F, Green CM, et al (2019) The non-canonical SMC protein SmcHD1 antagonises TAD formation and compartmentalisation on the inactive X chromosome. Nature communications 10: 30

Gendrel A-V, Attia M, Chen C-J, Diabangouaya P, Servant N, Barillot E & Heard E (2014) Developmental Dynamics and Disease Potential of Random Monoallelic Gene Expression. Dev Cell 28: 366–380

Hoki Y, Kimura N, Kanbayashi M, Amakawa Y, Ohhata T, Sasaki H & Sado T (2009) A proximal conserved repeat in the Xist gene is essential as a genomic element for X-inactivation in mouse. Development (Cambridge, England) 136: 139–46

Jansz N, Keniry A, Trussart M, Bildsoe H, Beck T, Tonks ID, Mould AW, Hickey P, Breslin K, Iminitoff M, et al (2018a) Smchd1 regulates long-range chromatin interactions on the inactive X chromosome and at Hox clusters. Nature structural & molecular biology 25: 766–777

Jansz N, Nesterova T, Keniry A, Iminitoff M, Hickey PF, Pintacuda G, Masui O, Kobelke S, Geoghegan N, Breslin KA, et al (2018b) Smchd1 Targeting to the Inactive X Is Dependent on the Xist-HnrnpK-PRC1 Pathway. Cell reports 25: 1912–1923.e9

Kalantry S & Magnuson T (2006) The Polycomb Group Protein EED Is Dispensable for the Initiation of Random X-Chromosome Inactivation. PLoS Genetics 2: e66

Kalantry S, Mills KC, Yee D, Otte AP, Panning B & Magnuson T (2006) The Polycomb group protein Eed protects the inactive X-chromosome from differentiation-induced reactivation. Nat Cell Biol 8: 195–202

Kato M, Takemoto K & Shinkai Y (2018) A somatic role for the histone methyltransferase Setdb1 in endogenous retrovirus silencing. Nat Commun 9: 1683

Keniry A, Gearing LJ, Jansz N, Liu J, Holik AZ, Hickey PF, Kinkel SA, Moore DL, Breslin K, Chen K, et al (2016) Setdb1-mediated H3K9 methylation is enriched on the inactive X and plays a role in its epigenetic silencing. Epigenetics & chromatin 9: 16

Kent WJ, Sugnet CW, Furey TS, Roskin KM, Pringle TH, Zahler AM & Haussler and D (2002) The Human Genome Browser at UCSC. Genome Res 12: 996–1006

Kohlmaier A, Savarese F, Lachner M, Martens J, Jenuwein T & Wutz A (2004) A Chromosomal Memory Triggered by Xist Regulates Histone Methylation in X Inactivation. PLoS Biology 2: e171

Kojima Y, Kaufman-Francis K, Studdert JB, Steiner KA, Power MD, Loebel D, Jones V, Hor A, de Alencastro G, Logan GJ, et al (2014) The Transcriptional and Functional Properties of Mouse Epiblast Stem Cells Resemble the Anterior Primitive Streak. Cell stem cell 14: 107–20

Leong HS, Chen K, Hu Y, Lee S, Corbin J, Pakusch M, Murphy JM, Majewski IJ, Smyth GK, Alexander WS, et al (2013a) Epigenetic Regulator Smchd1 Functions as a Tumor Suppressor. Cancer Res 73: 1591–1599

Leong HS, Chen K, Hu Y, Lee S, Corbin J, Pakusch M, Murphy JM, Majewski IJ, Smyth GK, Alexander WS, et al (2013b) Epigenetic Regulator Smchd1 Functions as a Tumor Suppressor. Cancer Research 73: 1591–1599

Li H & Durbin R (2009) Fast and accurate short read alignment with Burrows–Wheeler transform. Bioinformatics 25: 1754–1760

Li H, Handsaker B, Wysoker A, Fennell T, Ruan J, Homer N, Marth G, Abecasis G & Durbin R (2009) The Sequence Alignment/Map format and SAMtools. Bioinformatics 25: 2078–2079

Lyon MF (1961) Gene Action in the X-chromosome of the Mouse (Mus musculus L.). Nature 190: 372–373

Mak W, Nesterova TB, Napoles M de, Appanah R, Yamanaka S, Otte AP & Brockdorff N (2004) Reactivation of the Paternal X Chromosome in Early Mouse Embryos. Science 303: 666–669

Marahrens Y, Panning B, Dausman J, Strauss W & Jaenisch R (1997) Xist-deficient mice are defective in dosage compensation but not spermatogenesis. Genes & Development 11: 156–166

Matsui T, Leung D, Miyashita H, Maksakova IA, Miyachi H, Kimura H, Tachibana M, Lorincz MC & Shinkai Y (2010) Proviral silencing in embryonic stem cells requires the histone methyltransferase ESET. Nature 464: 927–931

Miura H, Takahashi S, Shibata T, Nagao K, Obuse C, Okumura K, Ogata M, Hiratani I & Takebayashi S (2020) Mapping replication timing domains genome wide in single mammalian cells with single-cell DNA replication sequencing. Nat Protoc 15: 4058–4100

Mohandas T, Sparkes R & Shapiro L (1981) Reactivation of an inactive human X chromosome: evidence for X inactivation by DNA methylation. Science 211: 393–396

Montgomery ND, Yee D, Chen A, Kalantry S, Chamberlain SJ, Otte AP & Magnuson T (2005) The Murine Polycomb Group Protein Eed Is Required for Global Histone H3 Lysine-27 Methylation. Current biology : CB 15: 942–7

Napoles M de, Mermoud JE, Wakao R, Tang YA, Endoh M, Appanah R, Nesterova TB, Silva J, Otte AP, Vidal M, et al (2004) Polycomb Group Proteins Ring1A/B Link Ubiquitylation of Histone H2A to Heritable Gene Silencing and X Inactivation. Developmental Cell 7: 663–676

Nesterova TB, Wei G, Coker H, Pintacuda G, Bowness JS, Zhang T, Almeida M, Bloechl B, Moindrot B, Carter EJ, et al (2019) Systematic allelic analysis defines the interplay of key pathways in X chromosome inactivation. Nature Communications 10: 3129

Nozawa R-SS, Nagao K, Igami K-TT, Shibata S, Shirai N, Nozaki N, Sado T, Kimura H & Obuse C (2013) Human inactive X chromosome is compacted through a PRC2-independent SMCHD1-HBiX1 pathway. Nature structural & molecular biology 20: 566–73

Okamoto I, Otte AP, Allis CD, Reinberg D & Heard E (2004) Epigenetic Dynamics of Imprinted X Inactivation During Early Mouse Development. Science 303: 644–649

Penny GD, Kay GF, Sheardown SA, Rastan S & Brockdorff N (1996) Requirement for Xist in X chromosome inactivation. Nature 379: 379131a0

Peters AHFM, O’Carroll D, Scherthan H, Mechtler K, Sauer S, Schöfer C, Weipoltshammer K, Pagani M, Lachner M, Kohlmaier A, et al (2001) Loss of the Suv39h Histone Methyltransferases Impairs Mammalian Heterochromatin and Genome Stability. Cell 107: 323–337

Plath K, Fang J, Mlynarczyk-Evans SK, Cao R, Worringer KA, Wang H, Cruz CC de la, Otte AP, Panning B & Zhang Y (2003) Role of histone H3 lysine 27 methylation in X inactivation. Science (New York, NY) 300: 131–5

Rens W, Wallduck MS, Lovell FL, Ferguson-Smith MA & Ferguson-Smith AC (2010) Epigenetic modifications on X chromosomes in marsupial and monotreme mammals and implications for evolution of dosage compensation. Proceedings of the National Academy of Sciences 107: 17657–17662

Sado T, Hoki Y & Sasaki H (2005) Tsix silences Xist through modification of chromatin structure. Developmental cell 9: 159–65

Sado T, Okano M, Li E & Sasaki H (2004) De novo DNA methylation is dispensable for the initiation and propagation of X chromosome inactivation. Development (Cambridge, England) 131: 975–82

Sado T & Sakaguchi T (2013) Species-specific differences in X chromosome inactivation in mammals. Reproduction (Cambridge, England) 146: R131–9

Sakakibara Y, Nagao K, Blewitt M, Sasaki H, Obuse C & Sado T (2018) Role of SmcHD1 in establishment of epigenetic states required for the maintenance of the X-inactivated state in mice. Development (Cambridge, England) 145

Sakata Y, Nagao K, Hoki Y, Sasaki H, Obuse C & Sado T (2017) Defects in dosage compensation impact global gene regulation in the mouse trophoblast. Development 144

Schotta G, Lachner M, Sarma K, Ebert A, Sengupta R, Reuter G, Reinberg D & Jenuwein T (2004) A silencing pathway to induce H3-K9 and H4-K20 trimethylation at constitutive heterochromatin. Gene Dev 18: 1251–1262

Silva J, Mak W, Zvetkova I, Appanah R, Nesterova TB, Webster Z, Peters A, Jenuwein T, Otte AP & Brockdorff N (2003) Establishment of Histone H3 Methylation on the Inactive X Chromosome Requires Transient Recruitment of Eed-Enx1 Polycomb Group Complexes. Developmental Cell 4: 481–495

Sugimoto M, Kondo M, Koga Y, Shiura H, Ikeda R, Hirose M, Ogura A, Murakami A, Yoshiki A, Chuva de Sousa Lopes SM, et al (2015) A Simple and Robust Method for Establishing Homogeneous Mouse Epiblast Stem Cell Lines by Wnt Inhibition. Stem Cell Reports 4: 744–757

Takahashi S, Miura H, Shibata T, Nagao K, Okumura K, Ogata M, Obuse C, Takebayashi S & Hiratani I (2019) Genome-wide stability of the DNA replication program in single mammalian cells. Nat Genet 51: 529–540

Tesar PJ, Chenoweth JG, Brook FA, Davies TJ, Evans EP, Mack DL, Gardner RL & McKay RDG (2007) New cell lines from mouse epiblast share defining features with human embryonic stem cells. Nature 448: 196–199

Trapnell C, Williams BA, Pertea G, Mortazavi A, Kwan G, Baren MJ van, Salzberg SL, Wold BJ & Pachter L (2010) Transcript assembly and quantification by RNA-Seq reveals unannotated transcripts and isoform switching during cell differentiation. Nat Biotechnol 28: 511–515

Wang C-YY, Jégu T, Chu H-PP, Oh HJ & Lee JT (2018) SMCHD1 Merges Chromosome Compartments and Assists Formation of Super-Structures on the Inactive X. Cell 174: 406–421.e25

Wang J, Mager J, Chen Y, Schneider E, Cross JC, Nagy A & Magnuson T (2001) Imprinted X inactivation maintained by a mouse Polycomb group gene. Nat Genet 28: ng574

Wutz A & Jaenisch R (2000) A Shift from Reversible to Irreversible X Inactivation Is Triggered during ES Cell Differentiation. Molecular Cell 5: 695–705

Zakharova IS, Shevchenko AI, Shilov AG, Nesterova TB, VandeBerg JL & Zakian SM (2011) Histone H3 trimethylation at lysine 9 marks the inactive metaphase X chromosome in the marsupial Monodelphis domestica. Chromosoma 120: 177–183

Żylicz JJ, Bousard A, Žumer K, Dossin F, Mohammad E, Rocha STT da, Schwalb B, Syx L, Dingli F, Loew D, et al (2019) The Implication of Early Chromatin Changes in X Chromosome Inactivation. Cell 176: 182–197.e23

